# POP-UP TCR: Prediction of Previously Unseen Paired TCR-pMHC

**DOI:** 10.1101/2023.09.28.560071

**Authors:** Nili Tickotsky

## Abstract

**Motivation:** T lymphocytes (T-cells) major role in adaptive immunity drives efforts to elucidate the mechanisms behind T-cell epitope recognition.

**Results:** We analyzed solved structures of T-cell receptors (TCRs) and their cognate epitopes and used the data to train a set of machine learning models, POP-UP TCR, that predict the binding of any peptide to any TCR, including peptide and TCR sequences that were not included in the training set. We address biological issues that should be considered in the design of machine learning models for TCR-peptide binding and suggest that models trained only on beta chains give satisfactory predictions. Finally, we apply our models to large data set of TCR repertoires from COVID-19 patients and find that TCRs from patients in severe/critical condition have significantly lower scores for binding SARS-coV-2 epitopes compared to TCRs from moderate patients (p-value <0.001).

**Availability and Implementation:** POP-Up TCR is available at: https://github.com/NiliTicko/POP-UP-TCR

## Introduction

Adaptive immune response requires that a T lymphocyte (T-cell) specifically recognizes, through its receptor (TCR), peptides presented on major histocompatibility complex (pMHC). Successful recognition leads to binding, that prompts conformational changes of the T-cell (Xu *et al*., 2020; Krogsgaard *et al*., 2003; Krogsgaard and Davis, 2005) followed by clonal expansion and acquisition of effector properties (Xu *et al*., 2020).

Successful prediction of bindings between TCRs and a specific peptide will enable the design of tumor-targeting T-cells, of peptides for vaccination and also the detection of autoimmune T cell clones.

The binding of a TCR to a pMHC is a pairing problem (Springer *et al*., 2021), where the goal is to predict whether a given pair of inputs X (a peptide) and Y (a TCR) would bind. Machine-learning solutions for this question vary according to the complexity level of the different pairing situations:

While at the more simple levels, the peptides are already seen in the training phase (Ostmeyer *et al*., 2019; Neuter *et al*., 2017; Meysman *et al*., 2018; Jokinen *et al*., 2020; Gielis *et al*., 2019; Fischer *et al*., 2020; Moris *et al*., 2020), the main challenge of TCR-peptide prediction models is the *de-novo* prediction of pairs in which neither TCR nor peptide has been in the train set (Moris *et al*., 2020). Current methods that deal with this situation report a higher than random, but generally less than satisfactory performance (Moris *et al*., 2020).

We hypothesized that a computational method that can perform *de-novo* prediction has to be based on general principles of TCR-peptide recognition that differentiate cognate pairs from pairs that do not functionally bind each other.

Our approach contains two stages: First we trained a machine-learning model based exclusively on structurally characterized TCR-pMHC complexes to predict, for each residue in complementary-determining region 3 (CDR3) of the beta chain of the TCR, whether it is likely to be in a physical contact with any peptide residue. The output of this structure-based model is a score for every pair of residues from CDR3 beta and the peptide. We then trained a second machine learning model on the scores given by the first model to thousands of sequences of TCR-peptide pairs. This model predicts whether the TCR as a whole is likely to actually bind the specific epitope peptide as a whole.

## Materials and Methods

We analyzed the structure of known 89 non-redundant complexes of TCRs bound to peptide-MHCs. This allowed the generalization of previous findings from a small number of TCR-complex structures(Xu *et al*., 2020; Singh *et al*., 2017) and the discovery of new principles that guide binding. Structures that contain both TCR chains and epitope peptides were obtained from the Protein Data Bank at the RCSB PDB website (http://www.rcsb.org/) (Berman *et al*., 2000) filtering by molecule type on the advanced search options. We used IMGT-numbered structure files from the Structural T-Cell Receptor Database (STCRDab)(Leem *et al*., 2017) at http://opig.stats.ox.ac.uk/webapps/stcrdab/. Using the coordinates in the ATOM line, TCR chains-peptide contacts were mapped defined as a distance of 6Å or less between respective c-beta atoms (c-alpha for glycine). BLOSUM62 sequence similarity matrix (R package ‘biostrings’)(Team, 2018) identified 89 non-redundant TCR alpha-beta-peptide structures.

We used these data (shown in the Supplementary material) to construct the features of the first, structure-based model (Figure 1).

**Figure 1:**
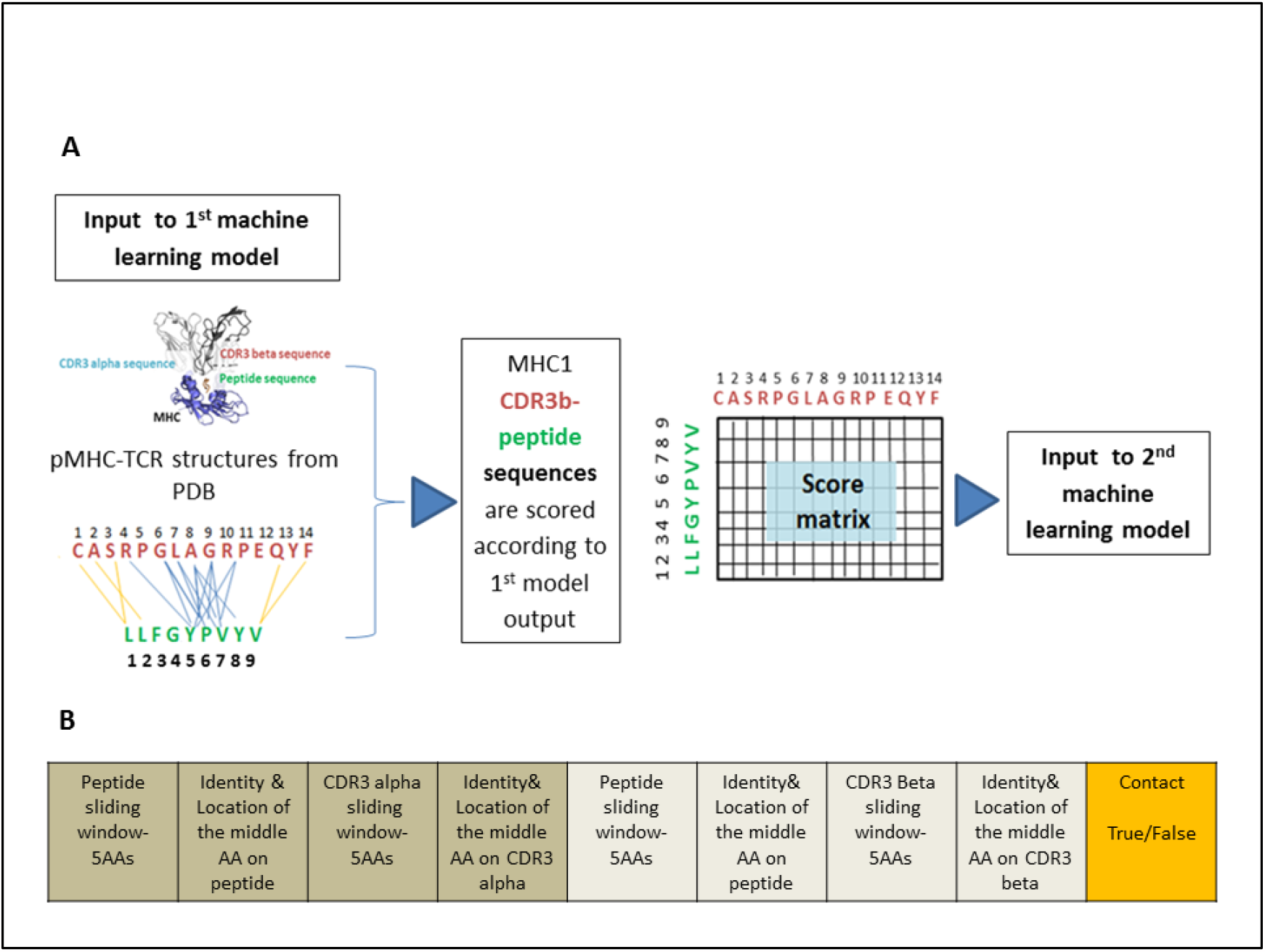
Prediction process workflow. **A.** The right side of the chart shows that all possible alpha-peptide and beta-peptide amino acid pairs serve as input to the first machine learning model that predicts whether they are likely to form a contact. Binding contacts are represented as blue lines, non-binding ones are in orange. For simplicity, not all non-binding pairs are shown. The output from the first model is a binding score for each pair of potential contact between the peptide and the CDR3. The predicted binding score of each beta-peptide amino acid pair is represented in a matrix that serves as input to the second machine learning model, which predicts whether or not this TCR recognizes and binds this peptide when presented on MHC1.**B**. Structure of the vectors for the 1^st^ model.

### Model for residue contact prediction

Based on the structural data, a set of features was composed per each TCR alpha-peptide pair, and per each TCR beta-peptide pair (Figure 2B). Each amino acid was represented as a one-hot vector of 21 numbers (20 possible amino acids and an additional stop codon) where all values were set to zeros except one index of the corresponding amino acid which was set to 1. To include all amino acids of the peptide and the CDR3 as given in the PDB database, the amino acids sequence of each was represented by a ‘sliding window’ of five amino acids, containing two amino acids adjacent to it on its N terminal side and two amino acids adjacent to it on its C terminal side. The one-hot vectors for all amino acids per ‘sliding window’ were joined. The use of the ‘sliding window’ is based on evidence that contact residues on the TCR are very specific both in the peptide residues they bind and those they allow in their vicinity. This specificity enables TCRs to discriminate between epitopes that differ even by a single oxygen atom (Mazza *et al*., 2007).

**Figure 2:**
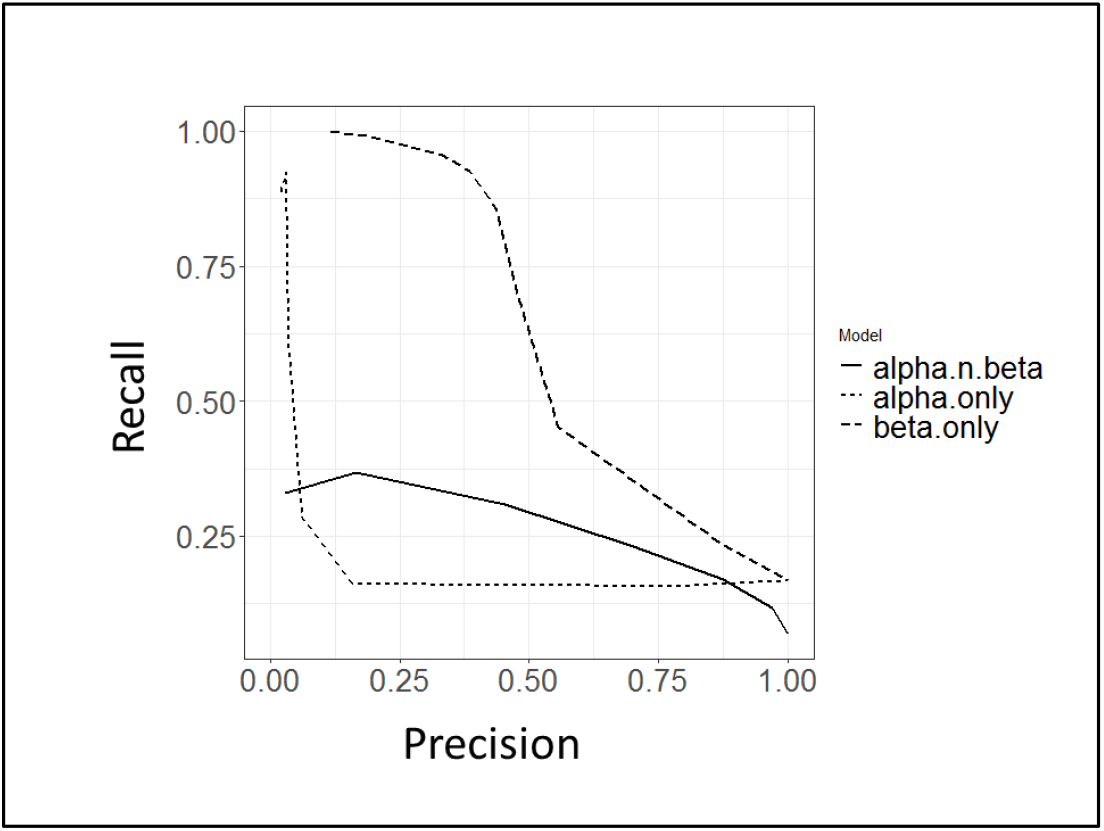
Precision-recall curves (PRC) of the performance of three machine learning models that learn from structural data. Each model predicts whether a given residue on a peptide and given residue on the CDR3 contact each other or not. A model based on the alpha CDR3 alone (dotted), a model based on the beta CDR3 alone (dashed), and a model that uses contacts from both chains without discriminating between them (solid).

Each vector contained a total of 28 features that described a pair of peptide-CDR3 residues and was labeled: ‘TRUE’ or ‘FALSE’ for a binding or a non-binding pair, respectively (Shown in Figure 1B). The features included:

1. A ‘sliding window’ for each CDR, five features per chain, ten in total.
2. The position number of the amino acid residue in the CDR (position 1 for the residue on the N-terminal end of the CDR) (See Figure 1 left hand side).
3. Two ‘sliding windows’ for the peptide: one for its contacts with CDR alpha and one for its contacts with CDR beta, five features per chain, ten in total.
4. A feature of the identity and a feature of the position of the amino acid of the peptide in contact with the TCR (position 1 for the residue on the N-terminal end of the peptide) (See Figure 1 left hand side).
5. A feature of the identity and a feature of the position of the amino acid of the TCR chain in contact with the peptide (position 1 for the residue on the N-terminal end of the peptide).

### Training and testing of residue contact prediction model

We trained three separate machine learning models to predict whether or not a given residue on a peptide and given residue on the CDR3 bind each other: A model based on the alpha CDR3 alone, a model based on the beta CDR3 alone, and a model based on both chains. (See Figure 1 for prediction process scheme). We used a 300-tree Random Forest (RF) algorithm implemented in R (R package ‘randomForest’). The performance of the models was assessed using a 5-folds cross-validation. Peptides in every fold were dissimilar to peptides in all other four folds.

### Performance analysis

Prediction scores for the test set were compared with the observed classifications, where a score (between 0 and 1) was considered as positive (i.e., binding) if it was above a certain cutoff. Precision and recall, as defined in the supplemental material were calculated for different cutoff values, and a precision-recall curve was created for each model.

### TCR - peptide binding sequence-based prediction model

#### Feature vector construction

The first model’s output was a score for the binding of each pair of amino acids-one on the TCR and one on the peptide. Each score was represented as an integer. All integers were joined and a padding integer of less than the minimal score was then added to the peptide vectors when it was necessary to complete to the maximum lengths as follows:

In the models with a single chain the number of possible pairs created by 15 peptide amino acids and 13 CDR amino acids is 15*13= 195. Models that use both the beta and alpha chains had twice that number, i.e., 290 features. 195 prediction scores per each position on the alpha and 195 prediction scores per each position on the beta chain-peptide contact from the first model were used as the set of features given to the second model (Figure 1). We expect that the distribution of predicted binding residues is different between TCR-peptide pairs that actually bind each other and pairs that do not and that these differences can be learned by the model.

Our training set for the second model was composed of a positive data set of 4,370 paired TCRαβ sequenced CDR3 segments with a cognate pMHC1 epitope retrieved from McPAS TCR database (http://friedmanlab.weizmann.ac.il/McPAS-TCR/), a manually curated database of T-cell receptor sequences (Tickotsky *et al*., 2017).

Redundancy reduction of peptide sequence was performed within the dataset itself and against the training set using the parameters that were used in the dataset construction (detailed in Supplementary material). Thus, no TCR-epitope peptide pair in the training set was redundant with any sequence in the test set or with any sequence in the PDB dataset.

Peptide sequences were divided into folds to accommodate for TCR cross-reactivity: This phenomenon, where a TCR may recognize multiple similar peptides (Krogsgaard *et al*., 2003; Gee *et al*., 2018; Birnbaum *et al*., 2014; Sewell, 2012; Hellman *et al*., 2019), can bias the performance of a machine learning model, if a peptide in the training set has a close homolog is in the test set. Indeed, the highest learning performance has been shown for epitopes that share some similarity with epitopes present in the training data(Moris *et al*., 2021). To address this issue we clustered the peptides in our dataset into folds by peptide similarity, where each fold contained all peptides that were similar to each other. Peptide inter-sequence similarity was measured by Levenshtein distance (LD)(Levenshtein VI, 1966), that sums the number of amino acid insertions/ deletions/substitutions . 232 unique peptide sequences were clustered into sixteen groups, each contained peptides that are similar by a total LD of four or less, which are equivalent to ∼40% similarity, with each peptide present in a single cluster. We set this strict similarity threshold based on evidence that a TCR can bind to peptides that are distinct at five or six positions from each other (Hausmann *et al*., 1999). The clusters were then joined to form five folds for training, verifying that no TCR-peptide pair in the training set is similar to a pair in the test set.

#### Negative dataset construction

TCR-epitope interaction databases contain only positive examples(Tickotsky *et al*., 2017; Shugay M, Bagaev DV, Zvyagin IV, Vroomans RM, Crawford JC, Dolton G, Komech EA, Sycheva AL, Koneva AE, Egorov ES, Eliseev AV, Van Dyk E, Dash P, Attaf M, Rius C, Ladell K, McLaren JE, Matthews KK, Clemens EB, Douek DC, Luciani F, van Baarle D, Kedzierska, 2017; Mahajan *et al*., 2018). We used the shuffling method (Moris *et al*., 2020), so our negative set was composed of the TCR alpha-beta chains from the training set randomly paired with a different peptide than that of the original pair. To reduce mislabeling of pairing TCRs with epitope-peptides to which they may cross-react, we clustered the epitope-peptides by sequence similarity before pairing them with the TCRs. As in the positive training set, we clustered the 232 unique epitope-peptides in the dataset into sixty-eight clusters, each cluster contained all peptides that were similar by at least 40% (LD = < 4). We then generated the random TCR-peptide pairs for the negative training and testing sets: each TCR’s alpha-beta CDR3 in the negative set was paired with epitope peptides from a different epitope cluster than that of the original pair in the train set, minimizing the chance for false negative results. Of the ∼250,000 pairs generated, we sampled 60,000 for our negative dataset.

#### Training and testing of binding protein prediction model

Training was performed using RF with 300 trees and five-fold cross validation. Performance was assessed with precision-recall curves, as described above. The test set included only peptides from clusters not found in the training set (termed “unseen epitopes” (Moris *et al*., 2021)). Again, we trained three separate machine learning models: A model based on data from the alpha CDR3 alone, a model based on the beta CDR3 alone, and a model based on both chains.

## Results

We analyzed 89 non-redundant X-ray crystal structures with both alpha and beta TCRs bound to MHC class 1 (MHC1) and MHC class 2 (MHC2) molecules from the PDB. The results of the structural data analyses are in the supplementary material.

### First model: Predicting residue-residue pairing

As our structural data showed that almost all peptide positions contact MHC, with almost no amino acids residues preferences (see Figure S1.A); we did not construct a model for the prediction of MHC-peptide contacts. Also, the differences between peptide contact distribution on the MHC1 and MHC2 (shown in Figure S1.A) and the paucity of MHC2-peptide binding data compared with the MHC1 led us to concentrate on MHC1-bound complexes.

Figure 2 shows the precision-recall curve (PRC) for predicting the bindings for each TCR chain. The model that was trained exclusively on the beta chain performed better than the other two.

### Second model: predicting peptide-TCR pairing

The residue pairing predictions of the first models described above classify each residue on the peptide for its potential to contact CDR3. However, residues that appear as putative CDR3 binding sites may sporadically occur also in peptides that do not actually bind CDR3. To predict whether the peptide and the CDR3, as wholes, actually bind each other we trained and tested another machine learning model, based on the prediction generated by the first classifier (scheme is shown in Figure 1). To train this machine, we relied on a positive set of 4370 TCR-peptide pairs that are known to bind each other and an assumed negative set of 60,000 pairs that are not known to bind each other (see Methods). We did not use, and did not have, any structural information for these pairs.

Figure 3 shows the precision-recall (PR) and receiver operating characteristic (ROC) curves from the three models tested. PR curve shows that the beta CDR3 model reached precision of 85% for recall level of 43%. For recall 12%, precision is 99%. ROC curve shows that the performance of the alpha-based model was similar to random (Area Under the Curve (AUC) of 0.474). The beta-based model and the model that uses both chains had an AUC of 0.780 and 0.748, respectively.

**Figure 3:**
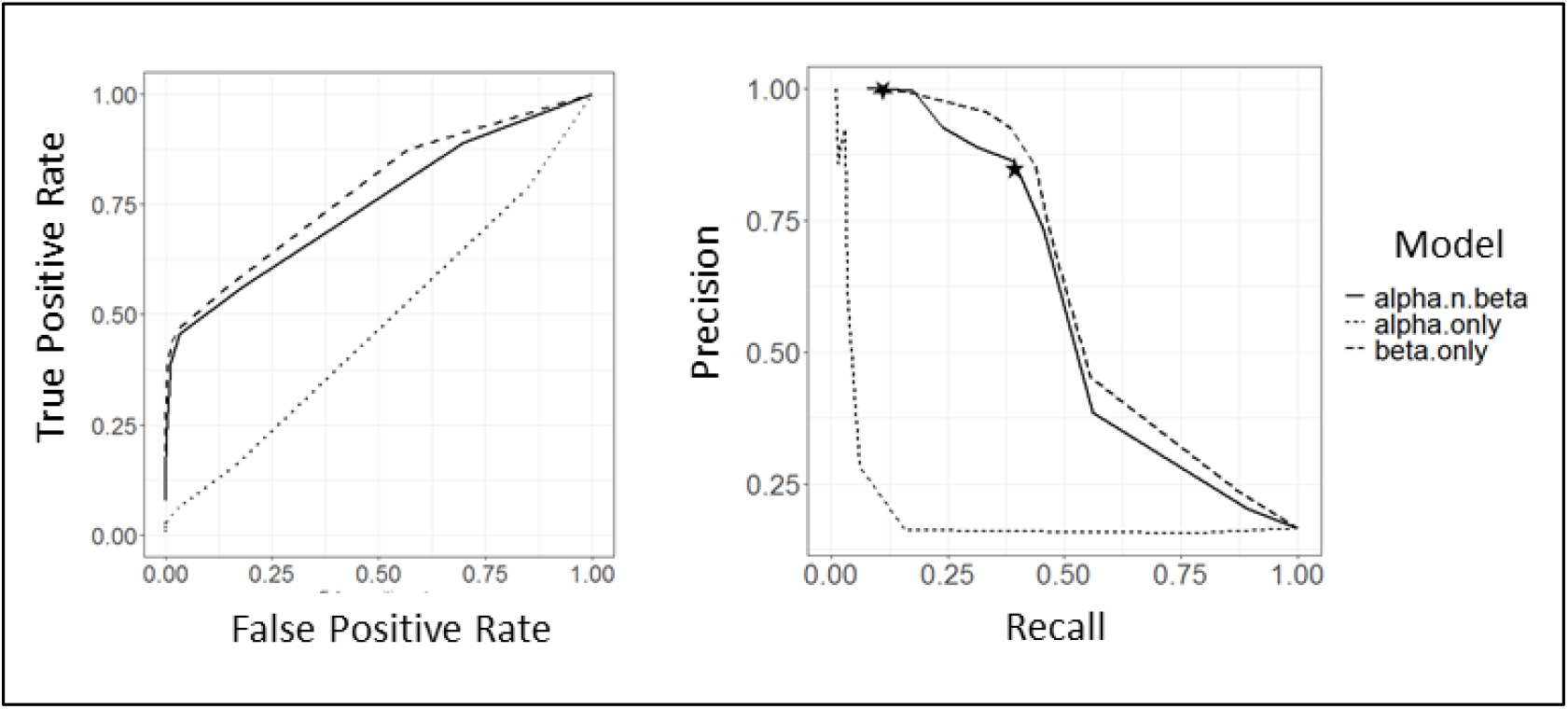
Precision-recall (PR) and receiver operating characteristic (ROC) curves for three separate machine learning models that that learn from sequence data and aim to predict whether a given peptide sequence and given CDR3 sequence bind each other or not. A model based on the alpha CDR3 alone, a model based on the beta CDR3 alone, and a model that is trained on both chains. The input to the prediction models is the sequence of a peptide and a sequence of a CDR3 beta. The output is a score for whether they bind each other or not. The test set is based on peptides new to the machine that were different by more than 40% from the peptides the machine had been trained on. For recall level of ∼43% the model correctly predicts 85% of the interacting pairs (point marked by a star). For recall 18%, precision is approximately 99% (point marked by a star).

### Benchmarking POP-Up TCR

For benchmarking, we tested the performance of POP-Up TCR prediction on the VDJdb dataset (Bagaev DV, Vroomans RMA, Samir J, Stervbo U, Rius C, Dolton G, Greenshields-Watson A, Attaf M, Egorov ES, Zvyagin IV, Babel N, Cole DK, Godkin AJ, Sewell AK, Kesmir C, Chudakov DM, Luciani F, 2019) as an independent test. We used a filtered dataset (Moris *et al*., 2020), as explained in the supplementary material.

As shown in Figure 4, for both modes of data down sampling the overall areas under the receiver operating characteristic (AUROC) were 0.5. This is very similar to results of other tools that tackle the unseen epitope problem, when benchmarked on complete different datasets (vs. on specific epitopes) (Moris *et al*., 2020). However, the precision-recall curves showed that only true binding CDR3beta-epitope pairs got the highest scores. This enables screening of true CDR3b-epitope pairs.

**Figure 4:**
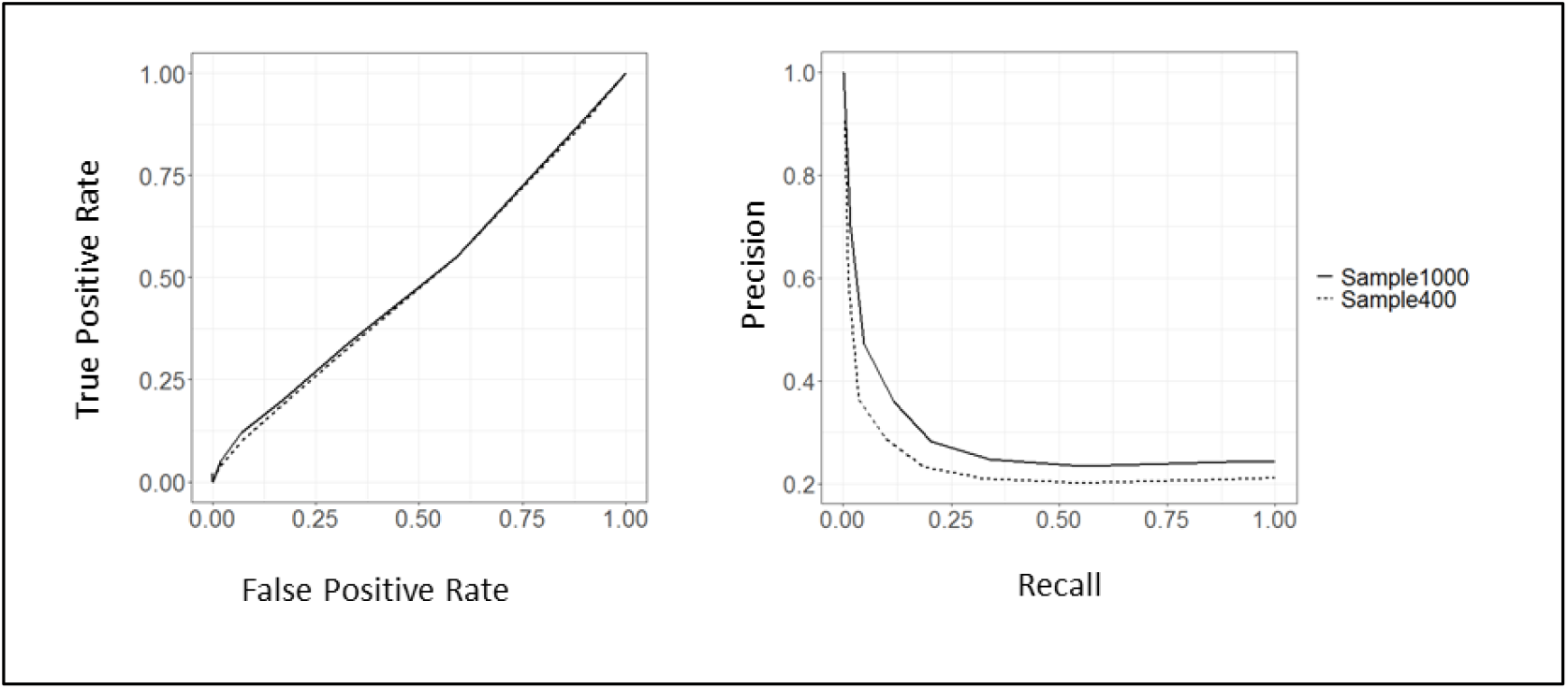
Performance of POP-UP TCR on an independent dataset, VDJdb (Bagaev DV, Vroomans RMA, Samir J, Stervbo U, Rius C, Dolton G, Greenshields-Watson A, Attaf M, Egorov ES, Zvyagin IV, Babel N, Cole DK, Godkin AJ, Sewell AK, Kesmir C, Chudakov DM, Luciani F, 2019).

### Prediction of cognate epitope-peptides for COVID-19 associated TCRs

We concluded that the best predictions on whether or not a given TCR will bind a given peptide can be obtained from the model trained only on beta chains, so we applied the exclusively beta chain-based models in both the first and second machine to TCRs retrieved from COVID-19 patients’ bronchoalveolar lavage fluid. Three patients had moderate disease, one had severe disease and five were defined critical cases(Liao *et al*., 2020). We tried to predict for the 40 most common SARS-Cov2 TCR CDR3 sequences found in these patients (these TCRs do not have a known cognate epitope) (Liao *et al*., 2020) their potential binding to epitopes derived from SARA-Cov2 proteins. We obtained 545 SARS-Cov2 epitopes From the Adaptive database (https://doi.org/10.21417/ADPT2020COVID) (Nolan *et al*., 2020). Of the 40 TCRs listed, twenty CDR3 sequences were from patients with moderate disease and twenty sequences were from patients who were in severe or critical condition. Each TCR’s beta CDR3 was submitted to the model with all 545 peptides, creating 21,800 prediction scores. The results of these predictions are shown in Figure 4.

TCRs from moderate COVID-19 patients have significantly stronger predictions for binding SARS-Cov2 peptide epitopes compared to TCRs from patients in severe/critical condition (Figure 5, Student’s t-test, p-value <0.001). Specifically, in the moderate group, 16 of the 20 TCRs had a least two peptides that got a prediction score above 0.3, with a total of 538 peptides that got this score. In the severe/critical group, only 9 of the TCR were predicted to bind five peptides with this score. In these patients, one of the TCRs got this score for 301 peptides. The difference between the TCRs in the two groups was their predicted ability to bind several epitopes, with TCRs in the moderate group displaying a greater cross reactivity than those in the severe/critical group.

**Figure 5.**
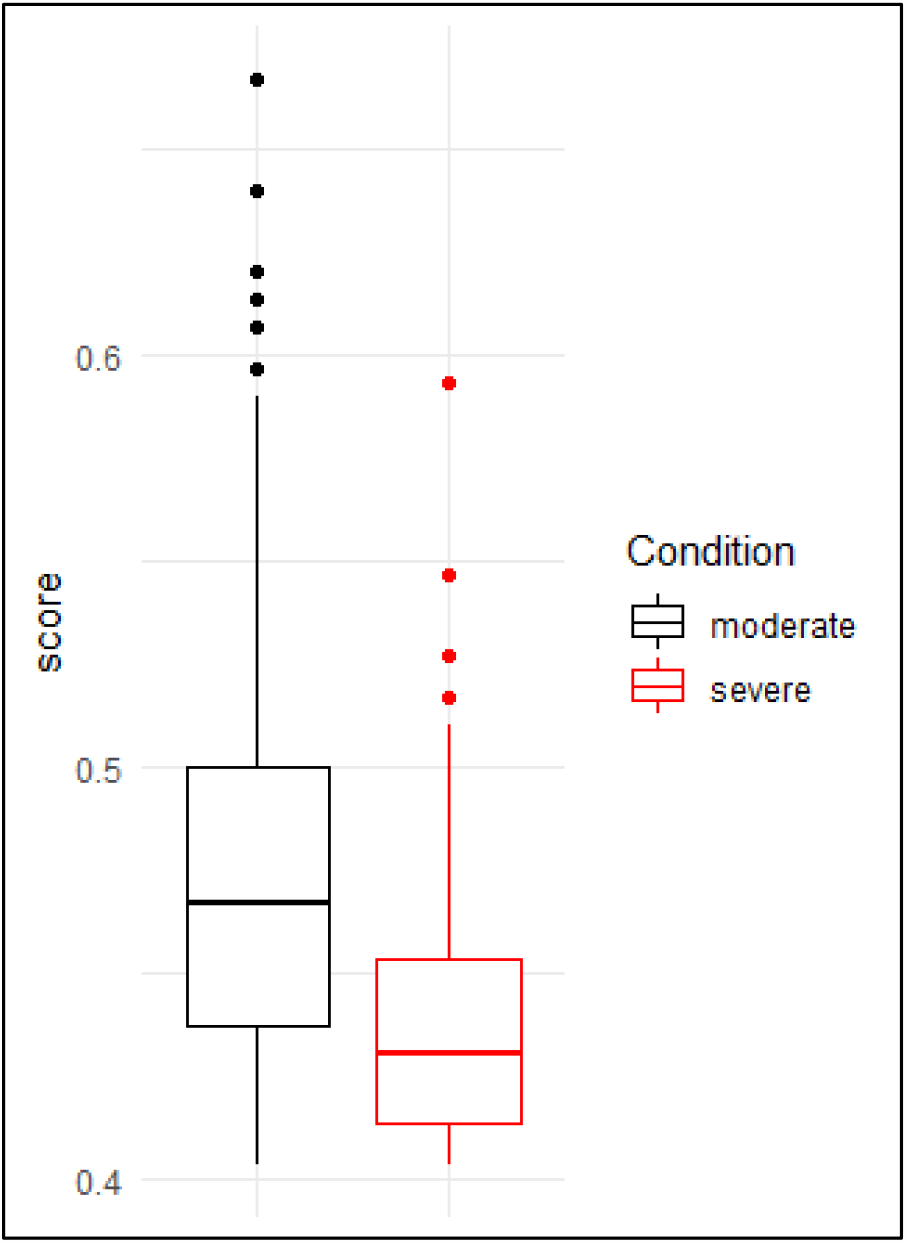
Model applications on the 40 most common SARS-Cov2 TCR beta CDR3 sequences that do not have a known cognate epitope(Liao *et al*., 2020). Twenty CDR3 sequences were from patients with moderate disease and twenty sequences were from patients who were in severe or critical condition. Each TCR’s beta CDR3 was submitted to the model with all possible 545 peptides, creating 21,800 prediction scores. TCRs from moderate COVID-19 patients have significantly stronger predictions for binding SARS-Cov2 peptide epitopes compared to TCRs from patients in severe/critical condition (Student’s t-test, p-value <0.001)

## Discussion

We believe the strength of our machine-learning approach stems from the fact that it places TCR-peptide binding in the context of the structural principles governing it.

Importantly, while the performance of the residue-residue contact prediction is far from perfect, it is significantly better than a random guess. Integrating the contact predictions into one score for the entire TCR and the entire peptide yielded strong predictions. We suggest this may mimic the nature of the binding process, where the affinity of the TCR to the peptide-MHC complex is a combination of much weaker residue-residue contacts. Some structural and chemical characteristics, such as specificity for a large hydrophobic residue, a charge, or a hydrogen bond donor, may induce conformational changes that lead to binding (Hausmann *et al*., 1999). Like us, recent studies have successfully used physicochemical (Moris *et al*., 2021; Ostmeyer *et al*., 2019; Karnaukhov *et al*., 2022) or biochemical (Beshnova *et al*., 2020) properties in both CDR and the peptide as features for machine learning models.

Others have demonstrated that different TCRs that bind to the same target often share sequence and structural features (Lanzarotti *et al*., 2019; Lin *et al*., 2021; Karnaukhov *et al*., 2022).

A major problem in TCR-peptide binding prediction is the absence of verified negative data (Moris *et al*., 2020). We suggest that pairing TCRs with peptides that belong to an entirely different cluster helped reduce the falsely labeled negative samples. We acknowledge, however, that with this design it is impossible to assess how well the method predicts the effect of small changes in a peptide. We refrained from using unpaired TCRs from healthy subjects in our negative data as such data probably includes pathology-related sequences (Madi *et al*., 2014).

Analyzing the experimentally solved TCR-peptide complexes, we found that the alpha and beta chains differ in their peptide binding preferences. Beta chains favor hydrophobic and anion-pi interactions, which contribute to binding (Zhu *et al*., 2021) and are relatively stable as they require less structural precision to obtain the same binding affinity of a peptide (Singh *et al*., 2017). The models that we trained on beta chain dataset alone outperformed the ones trained on the alpha chain or on mixed data sets composed of both alpha and beta chains sequences. This was true for both our models. The fact that we got good generic learning (i.e., of the unseen-epitope problem) with beta chain CDR3 data alone may be helpful for future deep learning models, as data on this chain is abundant in TCR-epitope databases.

A potential application of the model is to find the antigenic specificities of T cells of unknown specificity. We propose that using a computational model as the first step in such analyses may be a simpler and faster alternative that expands the available fitness landscape. We used our model to match TCRs from COVID-19 infected patients with potential cognate epitopes. Our prediction suggests that patients with severe disease may have fewer SARS-Cov2 TCRs than patients with mild disease.

This may be consistent with studies that found in antibodies that the potency of the immune response to SARS-Cov2 is predictive of survival (Garcia-Beltran *et al*., 2021).

## Acknowledgement

The author would like to thank Prof. Yanay Ofran for his great contribution in conceptualizing this study and his help with the manuscript.

