## Supplementary material for "POP-UP TCR: Prediction of Previously Unseen Paired TCR-pMHC"

**Structural data analyses:**

We first analyzed the structure of known 89 non-redundant complexes of TCRs bound to peptide-MHCs.

Epitope-binding TCR dataset: Structures that contain both TCR chains and epitope peptides were obtained from the Protein Data Bank at the RCSB PDB website (<http://www.rcsb.org/>) (Berman et al., 2000) filtering by molecule type on the advanced search options. We used IMGT-numbered structure files from the Structural T-Cell Receptor Database (STCRDab)(Leem, Oliveira, Krawczyk, & Deane, 2017) at <http://opig.stats.ox.ac.uk/webapps/stcrdab/>. Using the coordinates in the ATOM line, TCR chains-peptide contacts were mapped defined as a distance of 6Å or less between respective c-beta atoms (c-alpha for glycine). BLOSUM62 sequence similarity matrix (R package 'biostrings')(Team, 2018) identified 89 non-redundant TCR alpha-beta-peptide structures.

We analyzed 89 non-redundant X-ray crystal structures with both alpha and beta TCRs bound to MHC class 1 (MHC1) and MHC class 2 (MHC2) molecules from the PDB. We present here (Figure 2) the data gathered from these analyses on the contacts between residues on the TCR and residues on the pMHC complex. The architecture of TCR binding to peptides presented on MHC class 1 (pMHC1) is different than the binding of TCR to peptides presented on MHC class 2 (pMHC2)(Figure 2A). In pMHC1 the alpha CDR3 contacts the peptide on positions 1 to 7, counting from the N terminus, while the beta CDR3 contacts positions 3 to 10.

The average number of contacts formed with the TCR or MHC chain, in each position on the peptide is shown in Figure 2B. CDR3 on chain alpha forms most of its contacts with the 4^th^ and 5^th^ positions. CDR3 on the beta chain forms contacts that are more evenly distributed between the 5^th^and 8^th^ positions. MHC contacts are more evenly distributed, with preferences for more contracts as the N-terminus.

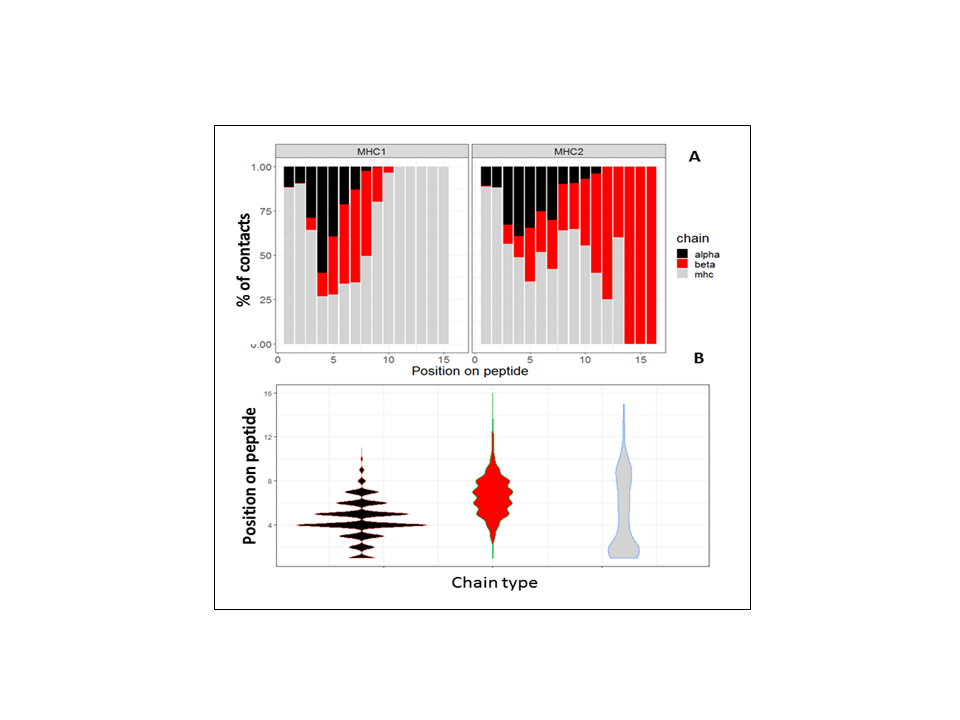

**Figure S1 Structural analyses results. A**. pMHC differ in the distribution contacts with alpha and beta CDR3s. In pMHC1, the alpha CDR3 contacts the peptide on its 1st to 7^th^ amino acids, while the beta CDR3 contacts it on its 3^th^ to 10^th^ amino acids, and the MHC1 contacts all the peptide's amino acids. In pMHC2, the alpha CDR3 contacts the peptide on its 1st to 12^th^ amino acids, while the beta CDR3 contacts it on its 3^th^ to 16^th^ amino acids, and the MHC2 contacts the peptide's 1^st^ to 13^th^ amino acids. **B.** Number of contacts on each position of the peptide: While most alpha CDR3 contacts are on the 4^th^ and 5^th^ positions, the beta CDR3 contacts are more evenly distributed between 5^th^, 6^th^ 7^th^ and 8^th^ positions. MHC contacts are quite evenly distributed, with the median on the 2^nd^ amino acid of the peptide.

**Peptide residue–chain residue preferences**

We compiled the log odds ratio of the observed frequency of each pair of amino acids over its expected frequency:

$$L_{X}x(i,j)={log}_{2}(\frac{P(i,j)}{P(i)P(j)})$$

Where the subscript x represents one of the three types of interfaces (peptide-CDR3 alpha, peptide-CDR3 beta, peptide-MHC), i and j are types of amino acids, P(i,j) is the probability of a contact between amino acids of type i and j in interfaces of type x, and P(i) and P(j) are the probability of occurrence for amino acids i and j, respectively, in the interfaces of type x. Here, to achieve maximal accuracy, a contact was defined as 5A between c-ceta atoms (c-alpha for glycine). Hence, the denominator describes the probability of a contact between i and j if the formation of contacts between i and j were random. For each chain we included only amino acid pairs that appeared at least five times, and from among those we considered as frequent only pairs that were at least 2 standard deviations from the mean likelihood L for that pair, as defined above.

Peptide residue–residue preferences differed between the chains, as can be seen in

Table S1. CDR3 beta- showed a preference for salt bridges, hydrophobic interactions and union-pi interaction. CDR3 alpha preferred proline, isoleucine, asparagine and glutamic acid on the peptide.

**Table S1** CDR- peptide amino acid residue preferences. CDR3 beta- showed a preference for salt bridges, hydrophobic interactions and union-pi interaction. CDR3 alpha preferred proline, isoleucine, asparagine and glutamic acid on the peptide.

| **Interaction type** | **Residue-residue contact preferences** | | |
| --- | --- | --- | --- |
|  | **Beta -peptide** | **Alpha- peptide** | **MHC- peptide** |
| **Hydrophobic** | E-L | I-M | V-T  L-V  A-D |
| **Salt bridge** | D-R | E-K |  |
| **Anion-pi** | W-D |  |  |
| **Aromatic-proline**  **(Proline on CDR)** |  | P-W  P-M  P-K |  |
| **H-bonds** |  | **N-S** | **Q-N**  **Q-Y**  **S-D** |

MHC chains- showed a preference for hydrophobic interactions and H-bonds. As hypothesized by Singh et al. (Singh et al., 2017), we found that at the MHC- peptide interface, salt bridges involve peptide termini and are located at the periphery of the interface, where desolvation penalties are reduced (Figure S2).

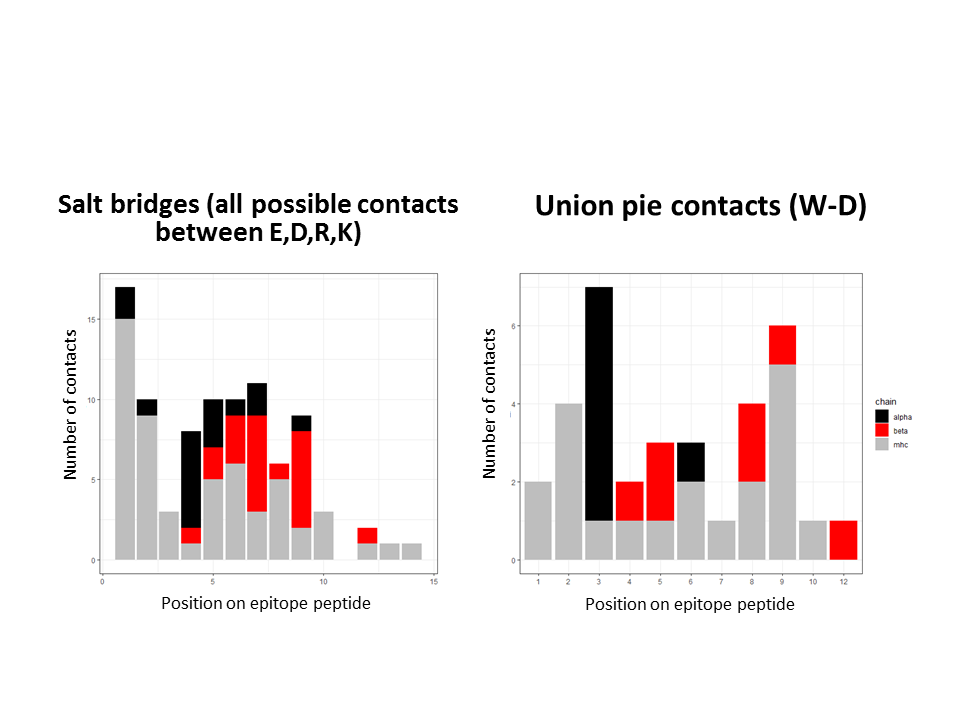

**Figure S2.** Peptide residue–residue preferences differed between the chains. CDR3 beta- showed a preference for salt bridges, hydrophobic interactions and union-pi interaction.

**Precision-Recall curves**

$$Precision=\frac{TP}{TP+FP}$$

$$Recall=\frac{TP}{TP+FN}$$

Where TP is the number of true positive predictions, FP is the number of false positives and FN is the number of false negatives. Expected performance at random was calculated as the fraction of positive examples from the total number of examples (which usually is equal to the precision value when recall equals 1).

**Benchmarking POP-Up TCR:**

For benchmarking, we tested the performance of POP-Up TCR prediction on the VDJdb dataset as an independent test. We used the filtered VDJdb dataset used by Morris et. al (Moris et al., 2020),that consisted of 14,188 CDR3 beta sequences. We took out CDR3 beta-peptide pairs that were in POP-UP TCR training data. As done by Moris et al., in order to avoid imbalanced learning of highly ubiquitous epitope peptides, we used either moderate or strong down sampling, and set a threshold of 1,000 and 400 examples per epitope respectively (reducing the beta chain dataset from 14,188 to 7645 and 6,362 sequence pairs respectively).
